# Evaluating the Utility of a Nanoscale Flow Cytometer for Detection of Surface Proteins on HIV and Extracellular Vesicles

**DOI:** 10.64898/2026.03.09.710614

**Authors:** Jonathan Burnie, Caroline Ouano, Vanessa Costa, Irene Castrosin, Catherine Hammond, Hannah Matthews, John Tigges, Kizzmekia S. Corbett-Helaire

## Abstract

**Background:** Flow virometry (FV) – the application of flow cytometry to viruses – has historically been hindered by the inability of cytometers to detect particles below ∼300 nm in size. However, advances in optics and fluidics have enabled cytometers primarily designed for cells to detect viruses and extracellular vesicles (EVs) through light scatter alone. In 2024, the CytoFLEX nano was released, marketed for the detection of particles as small as 40 nm; however, its performance has yet to be compared to a conventional instrument for FV.

**Methods:** FV was utilized to evaluate performance of the CytoFLEX nano and a conventional flow cytometer (CytoFLEX S). Instrument scatter sensitivity was assessed using NIST beads (40-400 nm), and virus stocks [human immunodeficiency virus (HIV), human coronaviruses (HCoV)-229E and HCoV-OC43]. For fluorescence analysis, HIV virions were stained with PE- and BV421-conjugated antibodies targeting virion incorporated proteins (CD38, CD44), individually and in combination. Finally, HIV stocks were labeled with antibodies against the envelope (Env) glycoprotein and tetraspanins (CD9, CD81) to assess EVs within virus preparations.

**Results:** Compared to the CytoFLEX S, the CytoFLEX nano exhibited substantially greater scatter sensitivity, reflected by up to 50-fold higher signal-to-noise ratio across NIST-traceable beads and virus samples. This enabled clearer resolution of smaller populations, including bead populations < 70 nm that were undetectable on the CytoFLEX S, as well as improved resolution across all viruses. While both instruments reliably detected stained proteins on HIV virions, the CytoFLEX nano revealed a distinct population of tetraspanin-positive EVs within HIV stocks that was undetected on the CytoFLEX S. Using GFP-tagged HIV, we identified Env^+^ particles lacking GFP, indicating the presence of Env on EVs.

**Conclusions:** The CytoFLEX nano exhibited markedly improved scatter sensitivity compared to the CytoFLEX S, improving detection of viruses and enabling detection of EV populations that were undetectable on the conventional instrument. While both platforms performed similarly for surface protein labeling, additional consideration of spectral overlap was needed with the CytoFLEX nano in multicolor experiments. These findings highlight that the complementary strengths of each platform can be utilized to more comprehensively characterize virus and EV populations, providing new opportunities to investigate nanoparticle heterogeneity.

## Background

Over the last two decades, the interest in applying flow cytometry to viruses - termed flow virometry (FV)-has become increasingly simple and feasible as cytometry instrumentation has improved (reviewed in^1–4^). FV leverages the quantitative, multiparametric, and high-throughput strengths of flow cytometry, enabling analysis of viruses across diverse applications, including diagnosis and monitoring of viral infections^5–7^, characterization of viral heterogeneity^8,9^, and for vaccine quality control^10,11^. Detecting viruses directly by light scatter through cytometry has traditionally been challenging since viruses typically fall within the range of instrument background (< 300 nm) on conventional instruments^3,4^. To address this, three common strategies have been routinely employed to permit FV: (1) specialized instrumentation with custom modifications to enhance small particle detection^5,12,13^, (2) fluorescent triggering to detect viruses with fluorescent labels (i.e., dyes/fusion proteins)^9,14–22^, and (3) bead-based methods that increase light scattering^7,23–26,26–28^.

Over the last decade, there has been an emergence of cytometers marketed for cellular analysis that also permit detection of viruses and EVs through light scattering alone^29–37^. Beckman Coulter CytoFLEX instruments (models S and LX) are one such instrument family. These instruments incorporate a series of advancements that facilitate visualization of small particles including avalanche photodiodes, wavelength-division multiplexing, violet scatter detection, and diode lasers to enhance light detection while reducing optical and electronic noise^29^. Using conventional CytoFLEX instruments, FV has previously been used as a tool to study murine leukemia virus^35–39^, HIV^8,40–44^ and SARS-CoV-2 pseudoviruses^45^, largely in the context of evaluating surface proteins without the need for magnetic beads, fluorescent tags, ultracentrifugation, or other auxiliary methods.

In 2024, a new small particle-dedicated instrument (CytoFLEX nano) was released marketed as having technical advancements which specifically facilitate the visualization of smaller particles such as EVs and viruses as small as 40 nm^46^. This enhanced sensitivity is achieved through the use of multiple scatter channels and extended laser dwell times, in addition to other instrument modifications. Notably, while the instrument has already begun being used to analyze the profiles of EVs^47–50^, it has yet to be applied to virus labelling. Furthermore, its potential to enable discrimination between vesicles and virions, an ongoing challenge in the field due to their overlapping physical and biochemical properties and nanoscale heterogeneity^28,51–53^, remains unexplored and presents an opportunity to better resolve and characterize these closely related nanoparticle populations.

Since surface protein labeling has been well established for HIV on the CytoFLEX S instrument^8,40–44^, we selected HIV as a model to compare staining on the conventional CytoFLEX S instrument with the small particle-dedicated nano instrument (CytoFLEX nano).

Herein, we compare the scatter and fluorescence sensitivity of the CytoFLEX nano to the CytoFLEX S for flow virometry applications including the detection of established virion incorporated proteins (CD38 and CD44) and EV markers. We find both instruments readily allow for the detection of single and dual antigens on the HIV surface, demonstrating comparable analytical rigor. However, the CytoFLEX nano offers a distinct advantage in detecting virions by light scatter with dramatically improved signal-to-noise ratios resulting in an enhanced dynamic range and limit of detection. The CytoFLEX nano also offered additional sensitivity for detecting EVs present within HIV preparations, which were unresolved by the CytoFLEX S. Leveraging this enhanced sensitivity, we probed the presence of tetraspanins as canonical EV markers in HIV stocks to assess the contribution of EVs in these samples. Despite the increased scatter sensitivity, when labeling both tetraspanins and the HIV envelope glycoprotein, we find that the CytoFLEX S enabled a more nuanced analysis of HIV-associated EVs due to an optical configuration which facilitated less spectral spillover. Taken together, we show that both instruments used in tandem can help expand our knowledge of individual virion surfaces.

## Methods

### Cell culture

All cells were grown in a 5% CO_2_ humidified incubator at 37 °C in complete media containing 10% FBS (GeminiBio, Cat# 100-106) and 1% penicillin/streptomycin (Life Technologies, Cat# 15140122). HEK293T cells (isolated from a female human embryonic kidney; ATCC, Cat# CRL-11268) were cultured in Dulbecco’s modified Eagle’s medium (DMEM, Gibco, Cat# 10313-021) while H9 cells (isolated from a 53-year-old male; BEI, Cat# ARP-87) were cultured in RPMI (Gibco, Cat#11875-093).

### Virus production

Virus production protocols were adapted from previously published methods^8,44^. For infection, H9 cells were pelleted and resuspended in 0.5 mL of undiluted HIV_IIIB_ virus, then incubated for 4 hours at 37 °C. Following infection, fresh media was added, and the cells were transferred to a T75 flask and cultured until the time of harvest (∼10-12 days). For virus production via transfection, HEK293T cells were seeded at a density of 0.6 × 10^5^ cells/mL in 6-well plates containing complete media. Transfection was performed once cells reached ∼70% confluence. A total of 1 μg of plasmid DNA was used for SG3ΔEnv (ARP-11051) pseudoviruses, and 1.5 μg for HIV iGFP (ARP-12455) and HIV Gag-iGFP JRFL (ARP-12456) viruses. All transfections were conducted using a 1:3 ratio of plasmid DNA (μg) to transfection reagent (μL) (PolyJet™ In Vitro Transfection Reagent, Fisher Scientific, Cat# 50-478-8). Culture supernatants containing virus were collected 48–72 hours post-transfection. All virus stocks were centrifuged at 300 × g for 5 minutes to remove cellular debris before being stored at −80 °C until use.

### Monoclonal Antibodies

The following antibodies were used for flow virometry staining: PE-conjugated antibodies from BD Biosciences, including CD38 (Cat# 555460), CD44 (Cat# 550989), CD81 (Cat# 555676), and CD9 (Cat# 312106), and from BioLegend, including CD103 (Cat# 350206), CD82 (Cat# 342104), and CD63 (Cat# 353004). The CD103 antibody was used as a negative staining control. BV421-conjugated antibodies were obtained from BD Biosciences for CD81 (Cat# 740079), CD9 (Cat# 743047), and CD63 (Cat# 740080), and from BioLegend for anti-human IgG Fc (Cat#M1310G05), CD38 (Cat# 397120) and CD44 (Cat# 338810). Anti-HIV-1 gp120 Monoclonal Antibody PGT128 (ARP-13352) was obtained from BEI Resources.

### Flow virometry staining and CytoFLEX S acquisition

Flow virometry was performed using a Beckman Coulter CytoFLEX S equipped with a standard optical configuration. All samples, including calibration beads, were run for 30 seconds on medium or fast before being recorded for 15 seconds at a sample flow rate of 10 μL/min using the tube loader. Direct and indirect staining protocols were adapted from previously published methods^8,42,44^. For all labeling experiments, cell-free viral supernatants were used at their undiluted titer, with an average particle concentration of ∼10^6-8^ particles/mL. For direct labeling, PE- or BV421-conjugated monoclonal antibodies were used at empirically determined concentrations (0.4– 1 μg/mL) and incubated with virus overnight at 4 °C in the dark. For indirect labeling, viruses were incubated overnight at 4 °C in the dark with 0.4 μg/mL of unlabeled PGT128 or isotype control antibody, followed by a 3-hour incubation with 0.5 μg/mL of PE-conjugated secondary antibody (anti-human PE, BioLegend, Cat# 410708). Antibody concentrations were optimized by titration to maximize signal-to-noise ratios (Fig. S1–S4). Following staining, samples were fixed in 2% PFA for 20 minutes. Prior to acquisition, all samples were further diluted in PBS to minimize coincidence.

Fluorescence calibration was performed using ViroCheck NanoParticle Reference Kit (Cat# V10425, Lot# 75221) or BD Quantibrite PE beads (Cat# 340495, Lot# 75221) where indicated. Light scatter calibration was conducted using NIST-traceable size standards (Thermo Fisher Scientific). Calibration was performed using FCM_PASS_ (https://nano.ccr.cancer.gov/fcmpass) as previously described^54^. Additional information on calibration procedures and MIFlowCyt-EV framework controls^55,56^ is provided in File S1. Uncalibrated data are shown in arbitrary units (a.u.). Where indicated, calibrated data are shown in estimates of Enveloped Virus (nm) calculated with FCM_PASS_ using the default refractive index (1.456). Flow virometry data were analyzed using FlowJo software (version 10.10.0) and are available on The NanoFlow Repository^57^.

### CytoFLEX nano flow virometry acquisition

Samples were stained as above for the CytoFLEX S and run on the CytoFLEX nano (Beckman Coulter, Brea, CA) with a standard optical configuration. The instrument was calibrated through the automated QC and Sensitivity Monitoring performed according to manufacturer’s specifications using the CytoFLEX nano Daily QC Scatterspheres, Fluorospheres, and Multi-intensity Fluorospheres (Beckman Coulter, Cat #C85323, #C86350, #C92889). QC values were used to set gains for experimental analysis and confirmed via reference standards (i.e., Exosome standards, fluorescent, Sigma-Aldrich, Cat# SAE0193). Violet-SSC1 (VSSC1-H) trigger was utilized for all data acquisition with a threshold ranging between 325-350. All samples, including calibration beads, were run at a flow rate of 1 μl/minute. Samples were acquired by time or total number of events as indicated in figure captions. Data analysis and scatter calibration were performed as above. Fluorescence calibration was performed using ViroCheck Nanoparticles. Signal-to-noise ratio were calculated by dividing the median scatter intensity of the particle population (virus or bead peak) by the median scatter intensity of the background population, as determined from violet side scatter histogram plots for both flow cytometers. All experiments presented in the main text figures were performed at least twice on both instruments.

## Results

### Comparison of scatter sensitivity for small particle detection

To begin comparing the small particle detection capabilities of the CytoFLEX S and CytoFLEX nano, we first analyzed NIST-traceable 100 nm beads on both instruments. Since this bead population is readily detected by violet side scatter on both platforms, it served as a baseline for comparing signal-to-noise ratios (S/N). By comparing the scatter signals between the bead population and background noise, we found that the signal-to-noise ratio was ∼735 on the CytoFLEX nano using the most sensitive violet side scatter channel (VSSCH-1), compared to ∼15 on the CytoFLEX S (Fig. 1A). This nearly 50-fold difference demonstrates that the CytoFLEX nano provides substantially greater separation between background noise and specific signal than the conventional instrument.

**Figure 1.**
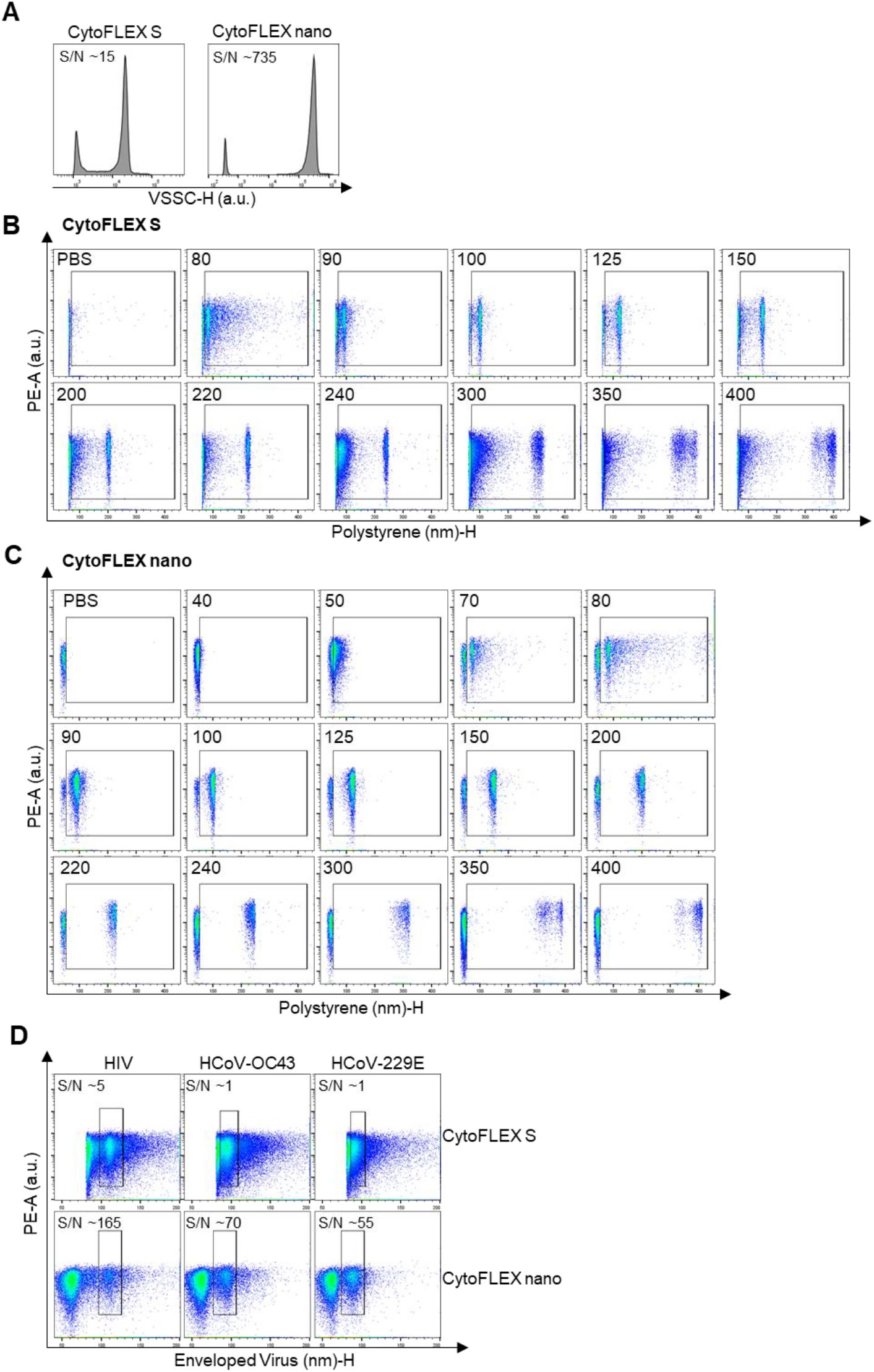
Comparison of calibration beads and viruses using the CytoFLEX S and CytoFLEX nano. (A) Detection of 100 nm polystyrene beads on the CytoFLEX S and CytoFLEX nano cytometers. Beads were resolved as distinct populations based on violet side scatter intensity (VSSC-H), with signal-to-noise ratios (S/N) indicated on the plots. Uncalibrated data are shown in arbitrary units (a.u.). (B) Pseudocolor dot plots for polystyrene beads ranging from 40 to 400 nm detected on the CytoFLEX S and (C) nano cytometers. Gates encompass the range of beads detected with scattering properties distinct from PBS. (D) Representative pseudocolor dot plots showing virus detection on both instruments, with S/N ratios indicated. Calibrated data are shown with enveloped virus sizing estimates generated through FCMPass using a refractive index of 1.456. Viral populations were identified by side scatter and are highlighted in the gated regions. Viral samples include HIVIIIB and human coronaviruses (HCoVs), HCoV-OC43 and HCoV-229E.

Next, to assess capabilities of the CytoFLEX nano for small particle detection across a larger scale, we ran a panel of NIST-traceable beads ranging from 40 to 400 nm (Fig. 1B-C). To compare the data on an equivalent scale, we calibrated the FCS files using FCMPass^54^ and displayed the data using polystyrene sizing estimates. With the calibrated scale, the beads showed close alignment with the anticipated size for both instruments (Fig. 1B-C). However, the 40, 50 and 70 nm bead populations were unable to be detected on the CytoFLEX S but were detectable on the CytoFLEX nano with the 70 nm population being fully separate from instrument background (Fig. 1C).

Since biological samples are known to refract light differently from polystyrene beads due to different refractive indices, we next sought to assess differences in scatter sensitivity across the two instruments using viruses. Since HIV has well established flow virometry protocols^40–44^, we used this virus as a reference standard. We additionally tested two human coronaviruses (HCoV-OC43 and HCoV-229E) to extend this analysis to a different class of RNA viruses which commonly circulate as seasonal pathogens. To this end, we ran unstained virus stocks for HIV, HCoV-OC43 and HCoV-229E on both instruments using violet side scatter for visualization (Fig. 1D). Although the CytoFLEX nano is also capable of evaluating scatter using the blue, yellow and red lasers, here analysis was limited to violet side scatter to allow for direct comparison to the CytoFLEX S. As expected, all three viruses – which range in size between ∼80-150 nm – were detected on both instruments by side scatter. While the HIV population was clearly resolved from background noise on both platforms, the coronaviruses showed less distinct separation. This suggests that HIV and CoV have differential light scattering properties despite similar sizes. In particular, HCoV-229E overlapped partially with background on the CytoFLEX S but was clearly distinguishable on the CytoFLEX nano (Fig. 1D).

Quantitatively, the CytoFLEX nano showed over a 50-fold increase in signal-to-noise ratio for the coronaviruses compared to the conventional CytoFLEX S instrument, similar to the signal-to-noise ratio observed with bead calibration (Fig. 1A), whereas the improvement for HIV was more modest (∼30-fold). The difference in S/N ratios across these viruses likely reflects intrinsic biological differences in their size, composition and light scattering properties. Attempts to lower the detection threshold on the CytoFLEX S resulted in an increased abort rate and introduced more events attributable to instrument noise and were therefore not pursued. Together, these data indicate that the CytoFLEX nano outperforms the conventional CytoFLEX S instrument in sensitivity for small particle detection, particularly for particles under 100 nm in size.

### Fluorescent labeling of cellular proteins on HIV virions

Given the importance of cellular proteins in HIV pathogenesis^58,59^, and the existence of established protocols for labeling these proteins on HIV^8,40,41,43,44^, we next sought to determine whether the enhanced sensitivity of the CytoFLEX nano provides any advantage over the CytoFLEX S for analyzing virion-incorporated proteins. To this end, we selected the virion-incorporated T-cell antigens CD38^44^ and CD44^60^ for labeling with PE-conjugated antibodies, since they have previously shown robust staining on virions from the H9 T-cell line using the CytoFLEX S^44^. To assess non-specific antibody labeling, two controls were used: media stained with anti-CD38 and anti-CD44 to measure background signal, and pseudoviruses (SG3 ΔEnv) produced in HEK293T cells lacking the proteins of interest to serve as antigen-specific negative controls.

When evaluating CD38 and CD44 staining on the CytoFLEX S, considerable levels of staining on the T-cell-derived virus (H9_IIIB_) were seen as expected, whereas negligible staining was seen on the media, pseudoviruses, and isotype control, (Fig. 2A). On the H9_IIIB_ virus, both cellular proteins were present on two distinct populations within the gated region, suggesting the presence of EVs with the proteins of interest, as seen previously^61^. As expected, the CytoFLEX nano provided a much larger dynamic range for scatter for visualization of smaller particles which are likely attributable to antibody background and labeled EVs (Fig. 2B, far left gate) that were not visible on the CytoFLEX S based on the threshold used (1100). Notably, the CytoFLEX nano provided largely similar fluorescent distributions of the labeled viral populations to those seen with the CytoFLEX S. Of note, when the same sample concentration and acquisition time were used on both instruments, the CytoFLEX S recorded substantially more events than the CytoFLEX nano (Fig. S5). This is likely due in part to the 10-fold higher acquisition rate of the CytoFLEX S compared to the CytoFLEX nano, but may also reflect increased instrument noise on the CytoFLEX S.

**Figure 2.**
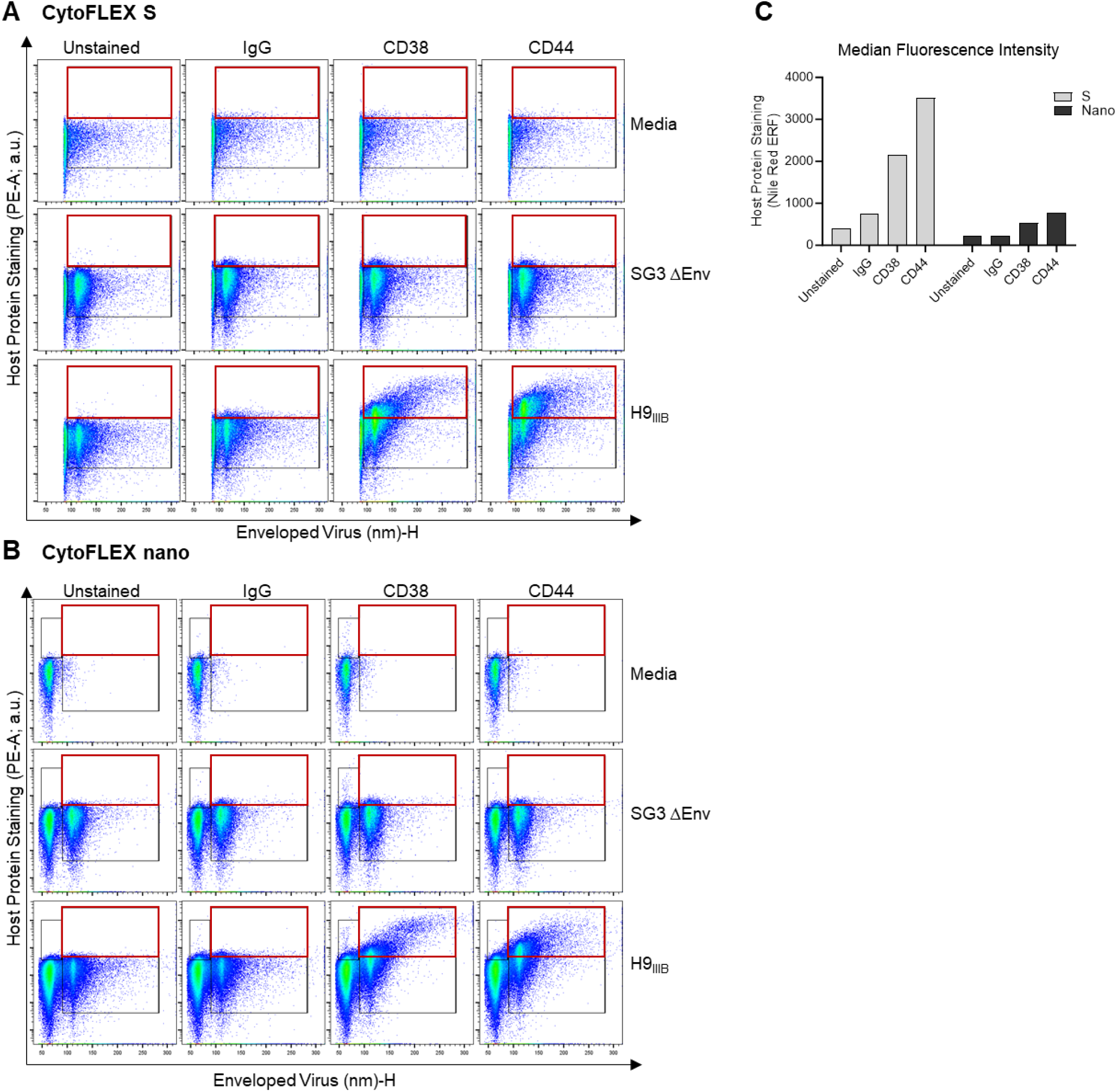
Detecting host proteins on HIV virions. (A-B) Presence of the cellular proteins CD38 and CD44 was assessed on HIVIIIB virions produced in H9 cells (H9IIIB) on the (A) CytoFLEX S or (B) CytoFLEX nano using PE-conjugated antibodies against the respective proteins. An isotype control antibody (IgG; targeting the irrelevant protein CD103) and unstained virus were used to assess nonspecific antibody binding and background fluorescence, respectively. HEK293T-derived pseudovirus, SG3 ΔEnv, is displayed as a negative virus control which lacks CD38 and CD44. Stained cell culture medium (DMEM) is shown to assess the contribution of antibodies in absence of virus to background fluorescence. In (A), the black gate is set based on the virus population as determined by scatter and is inclusive of the entire stained population, while the upper red gate denotes positive staining. In (B), the same gating strategy is shown as in (A), with an additional gate on the left of the x-axis set above instrument background fluorescence at the instrument threshold, which includes smaller particles including EVs and unbound antibody. (C) Median fluorescence intensity for the virus gate is shown for H9IIIB samples in calibrated equivalent reference fluorophore units (ERF) with Nile Red serving as the equivalent fluor for PE.

Although PE fluorescence is often reported in calibrated units of molecules of soluble fluorophore (MESF)^8,40^, MESF calibration beads were too large to be acquired on the CytoFLEX nano (∼3000 nm). Instead, when we ran Equivalent Reference Fluorophores (ERF) beads dedicated to small particles and compared fluorescence of the H9_IIIB_ samples, we found that the CytoFLEX S displayed higher PE values across all conditions (Fig. 2C). However, it should be mentioned that while the samples were run on the CytoFLEX S at the maximum PE gain, the CytoFLEX nano was run at a considerably lower gain based on the manufacturer’s QC guidelines. Thus, it is possible that increased fluorescent signal may have also been possible on the CytoFLEX nano.

Both instruments displayed relatively consistent mean fluorescence intensity across a virus dilution series over time (Fig. S6A–D). However, for unstained samples at lower virus concentrations, the CytoFLEX nano showed event counts that more closely aligned with the expected values based on serial dilution (Fig. S6E– H). The CytoFLEX S, however, exhibited increased variability unless stringent cleaning protocols (e.g., PBS or bleach washes) were applied between samples. These cleaning steps were unnecessary for the CytoFLEX nano since it has similar automated steps which occur in between each sample. However, these steps added considerable amounts of time to the acquisition of each sample compared to the CytoFLEX S (45 seconds on the CytoFLEX S vs. 4 minutes on the CytoFLEX nano).

### Dual labelling of cellular proteins on HIV

To assess the capacity of the nano to detect two antigens on the virus surface, we next stained CD38 and CD44 simultaneously on the same samples. For this purpose, we selected BV421 as a secondary fluorophore since it has previously shown to be effective for surface antigen labeling of HIV and MLV using the CytoFLEX S in FV^38,40^. Additionally, we tested both fluorophore-antibody pairings–CD38-PE with CD44-BV421 and CD38-BV421 with CD44-PE–to assess whether the choice of fluorophore affected staining patterns. We observed minimal staining on cell culture media and the negative control viruses, as expected for both instruments (Fig 3A; Fig S7). For H9_IIIB_, staining was seen for both CD38 and CD44 using PE- and BV421-labeled antibodies on the CytoFLEX S, as expected. While PE signals were comparable between the two instruments, the CytoFLEX nano exhibited higher BV421 signal relative to the CytoFLEX S, enabling clearer resolution of stained virus populations. These results demonstrate that the CytoFLEX nano can effectively detect dual antigen labeling on virus particles and may offer modest advantages over the CytoFLEX S in this application.

**Fig 3.**
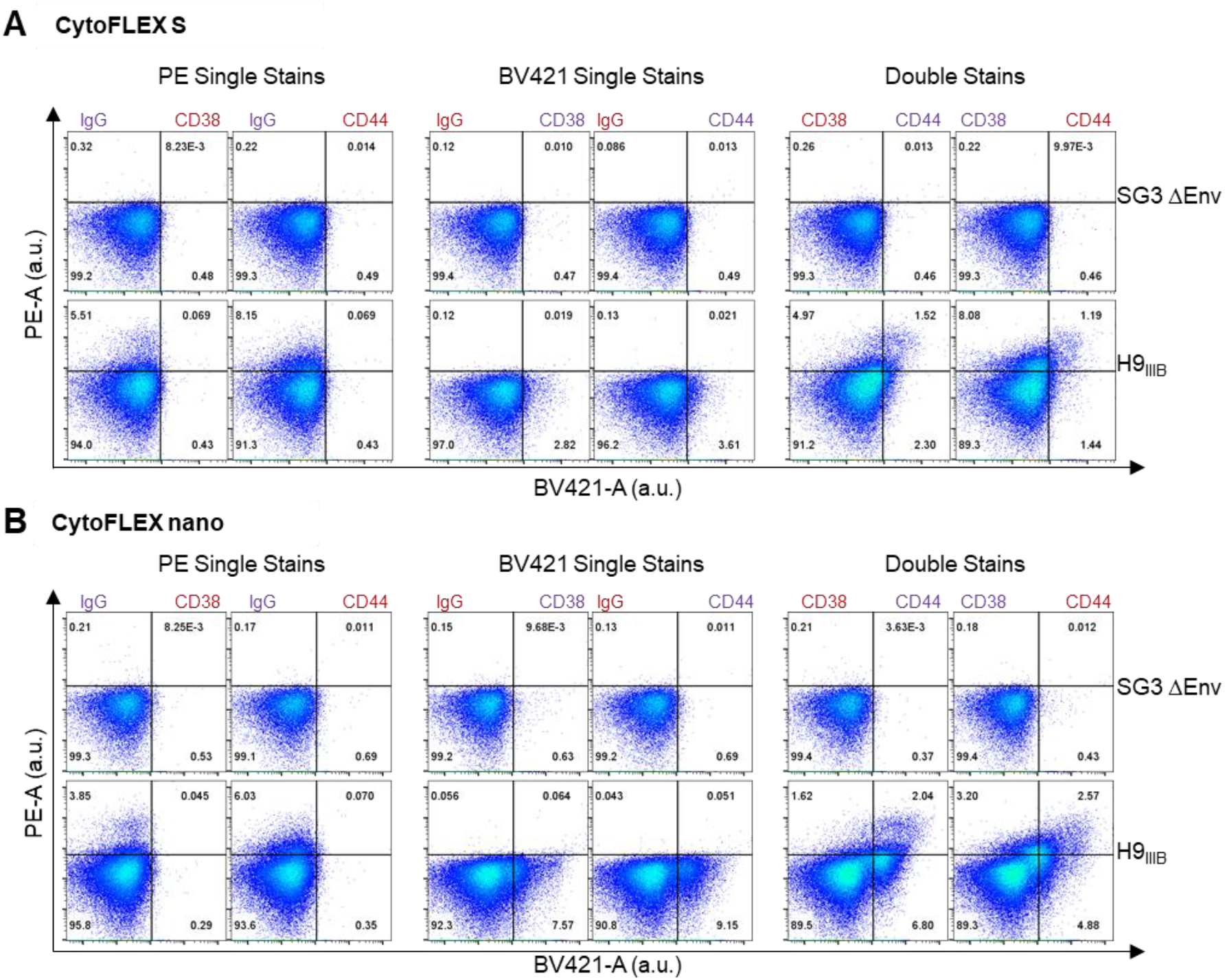
Comparing instrument utility for dual detection of host proteins on HIV. The presence of the cellular proteins CD38 and CD44 was assessed on H9IIIB virions on the (A) CytoFLEX S or (B) CytoFLEX nano using PE- and BV421-conjugated antibodies against the respective proteins. Isotype controls (IgG) were used to evaluate nonspecific staining for each channel. Red and purple labels represent PE- and BV421-labeled antibodies, respectively. The pseudovirus, SG3 ΔEnv is displayed as a virus control absent of CD38 and CD44. Gates are set based on staining with isotype controls on the SG3 Δenv virus stock.

### Labelling tetraspanins on HIV preparations

Since EVs are known to be present in virus preparations and share several surface markers with HIV^52,62,63^, we next sought to evaluate how the CytoFLEX nano could aid in uncovering the presence of EVs in virus preparations. Tetraspanins are a family of transmembrane proteins with a wide range of roles, including regulating cell morphology, motility, invasion and signaling^64,65^. Since tetraspanins such as CD9, CD63 and CD81 are widely recognized as canonical EV markers due to their consistent enrichment on EV membranes^66,67^, we sought to test for the presence of these markers on our virus stocks. CD82 was also added to the panel since it is an EV marker^68^ that was previously found to be highly abundant in HIV_IIIB_ preparations from H9 cells^44^.

When stained samples were analyzed using the CytoFLEX S, low levels of CD9 staining were observed in the media-alone condition, likely reflecting EVs from fetal bovine serum (Fig. 4A)^69,70^. Notably, EV marker profiles differed across the viruses: CD9 was detected on HEK293T-derived pseudoviruses (SG3 ΔEnv) but not on T-cell-derived (H9_IIIB_) viruses, where CD9 staining resembled that of the media control. In contrast, CD82 was abundant on H9-derived virions but absent on those from HEK cells, as to be expected based on published data^44^. CD63 and CD81 were detected on both virus types, with CD81 showing particularly strong staining on HEK-derived pseudoviruses.

**Fig 4.**
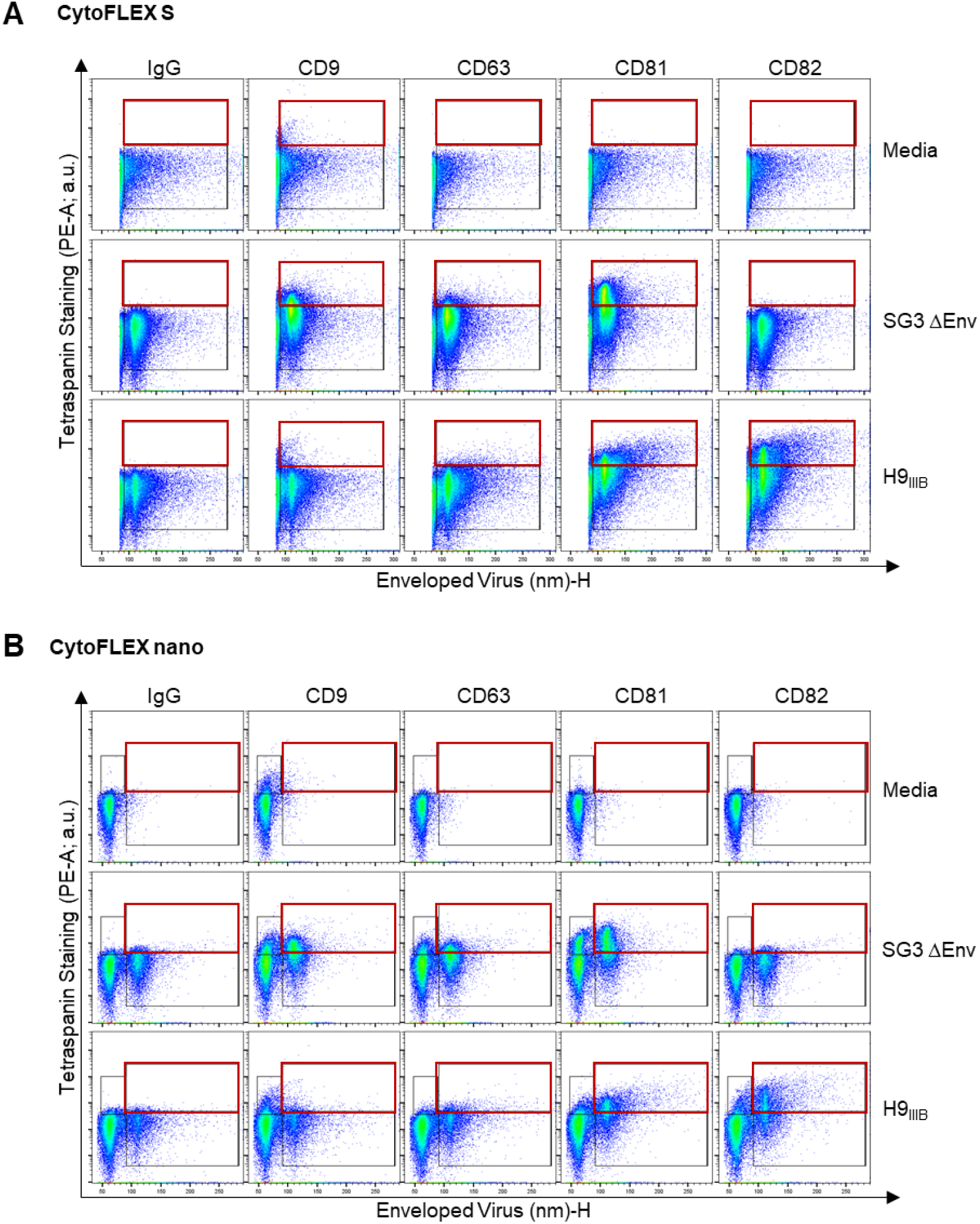
Detecting tetraspanins on HIV virions. Replication competent H9IIIB virions and SG3 ΔEnv pseudoviruses, were stained on the (A) CytoFLEX S or (B) CytoFLEX nano using PE-conjugated antibodies against tetraspanins (CD9, CD63, CD81, and CD82). An isotype control (IgG) is shown to assess nonspecific binding. A stained cell culture medium control is shown to assess contribution of antibodies alone (in absence of virus) to background fluorescence. In (A), the black gate is set based on the virus population as determined by scatter and is inclusive of the entire stained population, while the upper red gate denotes positive staining. In (B), the same gating strategy is shown as in (A), with an additional gate on the left of the x-axis set above instrument background fluorescence at the instrument threshold, which includes smaller particles including EVs and unbound antibody.

When the same samples were run on the CytoFLEX nano, a secondary scatter population was visualized that was completely unresolved on the CytoFLEX S. While the staining in the virus gate remained largely similar across instruments (Fig. 4B, right gate), staining in the EV gate (Fig. 4B, left gate) revealed that differences exist in the presence of EV populations within the HIV stocks. For instance, while CD63 showed relatively low levels of staining in the virus gate for both viruses, the presence of CD63^+^ EVs were detected in the pseudovirus sample which were absent in the replication-competent virus sample. Strikingly, a large CD81^+^ population of EVs were present in the pseudovirus sample that was less abundant in the replication-competent virus. Overall, these findings demonstrate that the CytoFLEX nano enables improved resolution of EV populations within virus preparations that were undetectable using the CytoFLEX S instrument alone.

### Evaluating the presence of EVs in HIV preparations

While tetraspanins are known EV markers, they are not exclusive to EVs and can also be present on replication-competent virions^71,72^. To begin addressing the overlap of viruses and EVs in virus preparations, we chose to stain the HIV envelope glycoprotein in tandem with tetraspanins on HIV stocks for testing on both cytometers. For this purpose, we utilized a GFP-expressing recombinant HIV construct which has GFP inserted between the structural proteins capsid and matrix^73^. Thus, the presence of GFP fluorescence in this system is indicative of the viral capsid protein which is more likely to be abundant in virions compared to EVs.

To begin, we stained two virus stocks with the broadly neutralizing anti-Env antibody PGT128^74^, using a previously described indirect staining protocol^42^. One virus stock contained a subtype B, R5-tropic envelope from the HIV JR FL strain (iGFP JR FL), while the other was a control virus lacking Env (iGFP ΔEnv). When run on the CytoFLEX S, PGT128 provided robust labeling on the Env^+^ virus but not on the ΔEnv virus, as expected. Notably, while the majority of staining was seen on GFP^+^ particles (i.e., virus population), a smaller population of Env^+^ events were present that did not display GFP, suggesting that these are EVs with HIV Env (Fig. 5A). This was in line with results from Arakelyan *et. al*., wherein a magnetic nanoparticle-based FV approach was utilized^28^.

**Fig 5.**
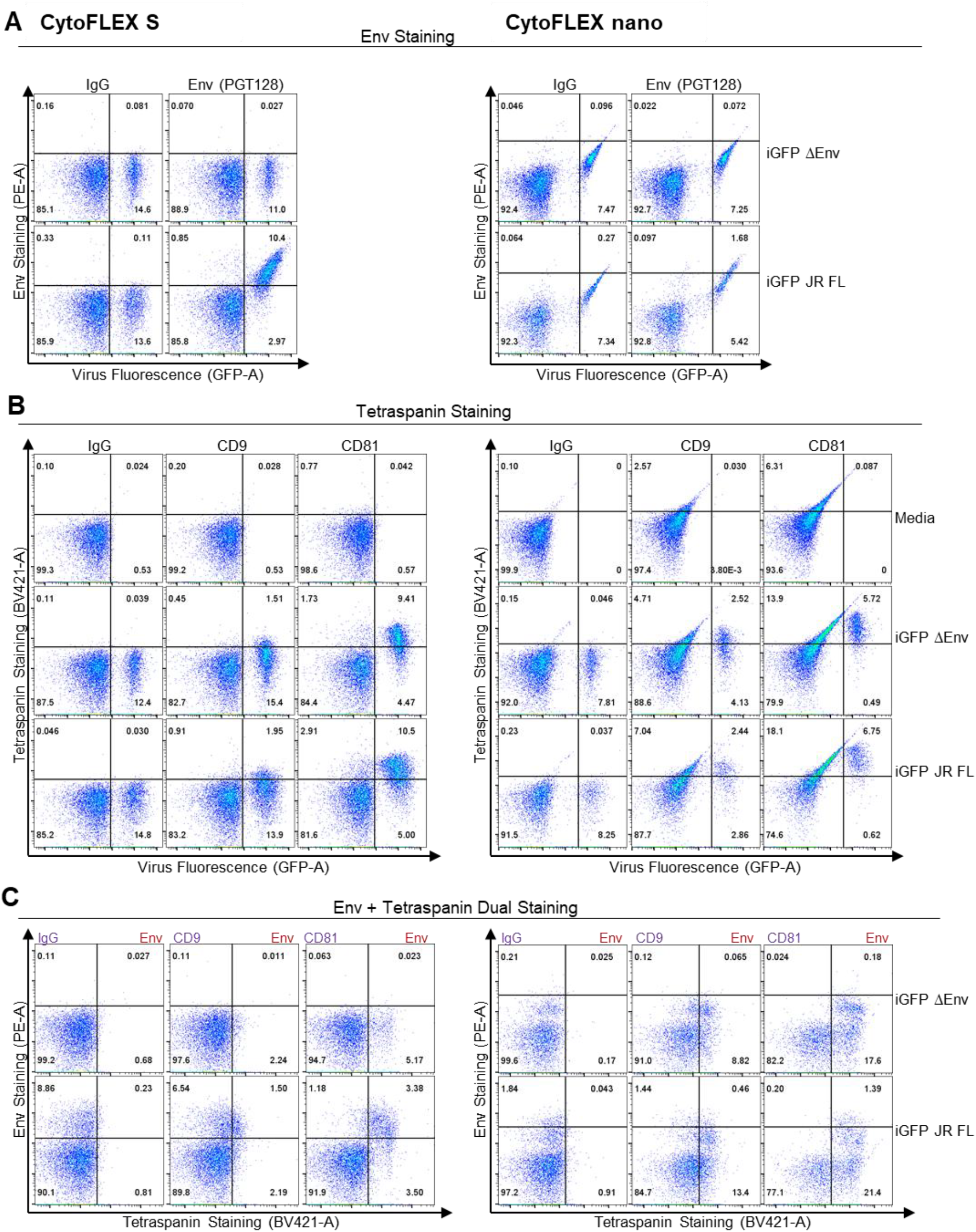
Evaluating presence of extracellular vesicles in HIV preparations. (A) Envelope glycoprotein staining was performed on the GFP-tagged infectious molecular clone iGFP, with or without (ΔEnv), the JR FL envelope. Samples were stained using either an isotype control or the anti-Env antibody PGT128, followed by detection with a PE-conjugated secondary antibody. Data were acquired on the CytoFLEX S (left panels) and the CytoFLEX nano (right panels). (B) Tetraspanin (CD9 and CD81) staining on viruses was performed using BV421-conjugated antibodies. Cell culture medium controls denote the contribution of antibodies alone to fluorescent signal. (C) Dual staining of HIV envelope glycoprotein and tetraspanins on HIV iGFP stocks. Gates are set based on staining with isotype controls on the ΔEnv virus stock. All fluorescence is displayed in arbitrary units.

Unfortunately, because the CytoFLEX nano has a co-linear laser detection configuration, it has a higher propensity for spectral overlap than the CytoFLEX S. As a result, when the same samples were run on the CytoFLEX nano, all GFP^+^ events were also positive in the PE channel without compensation, impacting detection of Env which was also detected in the PE channel (Fig. 5A). Despite this, a clear difference in staining was still seen on the CytoFLEX nano for Env staining on the Env^+^ and ΔEnv viruses.

Next, we performed tetraspanin staining on the same GFP-tagged virus stocks, focusing on CD9 and CD81 as representative tetraspanins of low and high abundance on HEK cell-derived viruses, respectively (Fig. 4). When analyzed on the CytoFLEX S, two distinct positive populations were observed for both CD9 and CD81: one GFP^+^ and one GFP^-^(Fig. 5B). This finding aligns with our prior CytoFLEX nano results, which revealed a distinct EV population within HIV preparations expressing these tetraspanins (Fig. 4B). While the same populations were present on the CytoFLEX nano, the enhanced BV421 signal from the CytoFLEX nano resulted in a distribution that was different from the S, indicative of spectral spillover.

Finally, we stained both Env and tetraspanins on the virus preparations. On the CytoFLEX S, clear Env^+^ and tetraspanin-positive populations were present on the JR FL Env virus but not the ΔEnv control, as expected (Fig. 5C). For both CD9^-^ and CD81^-^ labeled samples we saw three distinct populations within HIV stocks: (1) virions bearing Env and tetraspanins, (2) virions with Env that lack tetraspanins, and (3) EVs bearing tetraspanins but lacking Env. On the CytoFLEX nano, while the same populations were seen, spillover between GFP from the viruses and PE from Env staining resulted in all GFP^+^ viruses being detected as PE positive. Thus, while both instruments allow for multicolor labeling, compensation is more likely to be required on the CytoFLEX nano, even for color combinations that did not require compensation on the S.

## Discussion

While detection of unlabeled virions was previously challenging on conventional cytometers due to instrument limitations^1,3,4^, newer instruments have made flow virometry an increasingly accessible tool for single-virion characterization. Here, we demonstrated that while conventional cytometers like the CytoFLEX S provide strong utility for evaluating surface proteins on HIV, small particle-dedicated instruments, such as the CytoFLEX nano, offer clear advantages for analyzing viruses and EVs under 120 nm. The most notable difference between the instruments was scatter sensitivity, where the CytoFLEX nano provided dramatically improved signal-to-noise ratios and enabled detection of populations entirely undetectable on the CytoFLEX S. This enhanced resolution is particularly relevant given the growing interest in virus-associated EVs and their potential roles in infection^28,75–77^.

While the CytoFLEX nano performed relatively comparably to the CytoFLEX S for labeling surface antigens such as CD38 and CD44 on HIV virions, it did not exhibit notable improvements in fluorescence sensitivity in the PE channel. This is notable, as PE is commonly used for flow virometry^8,36–44^ because its intrinsic properties (size and brightness), together with the availability of reference materials, make it well suited for quantitative labeling of surface proteins^78,79^.

Importantly, although CD38 and CD44 were abundant in the virions analyzed here, many relevant viral proteins are expressed at much lower levels^37,58,59,80^. Thus, enhanced fluorescence sensitivity remains a key need for future instruments to enable reliable detection of low abundance viral surface proteins^1^. Additionally, we observed differences in virus staining profiles when labeling the same antigen with PE versus BV421, despite both being considered bright fluorophores in cellular flow cytometry. This highlights the potential to further optimize brighter fluorophores specifically engineered for small particle applications to maximize detection of low abundance antigens.

The ability to distinguish virions from EVs remains a critical hurdle in the field due to the overlapping size range (∼100-120 nm), biogenesis pathways, and shared surface markers^52,63^. Leveraging enhanced scatter sensitivity of the CytoFLEX nano, we detected EVs in HIV stocks with scatter and antigenic signatures distinct from virions. We hypothesize these EVs are most likely to be exosomes (30–150 nm) and microvesicles (50–1000 nm) due to their size ranges which overlap with those of HIV^81–83^. However, as EV biology and cytometric technologies continue to advance, future work should entail more detailed discrimination of the specific populations of EVs that exist within virus preparations. Previously, Arakelyan *et al*. showed that Env was present on HIV-associated EVs by using a magnetic nanoparticle-based flow virometry approach^28^. Our result shows that improvements in flow cytometry instrumentation can facilitate more precise visualization of Env on EVs without the requirement of magnetic nanoparticles. In this study, we also utilized an infectious molecular clone with structural proteins that are GFP-tagged^73^ to distinguish between virions and EVs. However, developing protocols that enable internal staining of capsid proteins in wild-type virions would further enhance the power of small particle cytometry for characterizing EV content in viral preparations.

Despite the increased scatter sensitivity of the CytoFLEX nano, we found that the conventional instrument remained better suited in some key areas for flow virometry. For instance, the altered optical figuration on the CytoFLEX nano resulted in increased spectral overlap, particularly between GFP and PE channels and GFP and BV421 channels, which impacted dual labeling experiments targeting the HIV Env and tetraspanins. In contrast, the CytoFLEX S provided more distinct spectral separation for multicolor analysis. Moreover, while the CytoFLEX nano allowed for increased sensitivity, this is partially achieved by a ten-fold reduction in the sample flow rate (1 µl/min vs 10 µl/min) and automated cleaning protocols which are run between each sample. These features increase overall experimental acquisition length, reducing the capacity for high throughput applications. Thus, rather than a replacement for conventional instruments, the CytoFLEX nano can complement existing platforms with its distinct strengths for small particle analysis. Of note, although many FV studies omit wash steps to remove unbound antibodies^8,37–44^, the increased sensitivity of the CytoFLEX nano allowed for visualization of antibody aggregates in our flow plots, highlighting the need for careful antibody titration to minimize non-specific signal.

While virus sorting was not performed in this study, integrating it with the enhanced scatter sensitivity of the CytoFLEX nano could enable more detailed analysis of viral heterogeneity and the high proportion of defective particles within virus stocks – a feature especially prominent in HIV, where most virions produced during infection are non-infectious^84^.

## Conclusions

In this study, we demonstrate that the CytoFLEX nano is comparable to the CytoFLEX S for phenotyping HIV surface proteins using established surface staining protocols. However, the CytoFLEX nano exhibited markedly improved sensitivity for detecting small particles by light scatter. This improved light scatter sensitivity revealed vesicle populations within HIV stocks that were undetectable on the CytoFLEX S, offering new opportunities to investigate virus-vesicle heterogeneity. However, spectral spillover on the CytoFLEX nano limits its suitability for complex multicolor analyses, and lengthy sample loading times reduce its utility for high-throughput applications. These results highlight the value of leveraging both platforms to achieve a fuller and more nuanced understanding of virus and vesicle populations.

## List of abbreviations

a.u.: arbitrary units
PE: Phycoerythrin
MESF: Molecules of Equivalent Soluble Fluorochrome
HIV: Human Immunodeficiency Virus
HCoV: Human Coronavirus
FBS: Fetal Bovine Serum
FV: Flow Virometry
ERF: Equivalent Reference Fluorophores
EV: Extracellular Vesicle
Env: Envelope Glycoprotein
MFI: Median Fluorescence Intensity
SSC: Side Scatter
S/N: Signal-to-noise ratio.

## Declarations

### Ethics approval and consent to participate

Not applicable.

### Consent for publication

All authors approved the manuscript for publication.

### Availability of data and materials

Source data supporting the findings of this study are available within this paper and its supplementary files. All FCS files are available on nano flow repository ID# 4851013737. Additional source data and materials inquiries should be directed to the corresponding author, Kizzmekia Corbett-Helaire. Materials will be made available after completion of a Material Transfer Agreement (MTA). If the material was obtained under use restriction, the inquiry will be forwarded to appropriate party.

### Author Contributions

Conceptualization – JB, VC, JT; Investigation – JB, CO, VC, IC, GH, HM, CH; Resources – JT, KSC; Supervision – JT, KSC; Writing (original draft) – JB; Writing (review & editing) – all authors.

## Acknowledgments

We thank Fidan Baycora for administrative support and Emily Hobbs for grant support. Reagents were obtained through the BEI Resources, National Institutes of Allergy and Infectious Diseases, National Institutes of Health: H9 Cells, ARP-87; HIV-1 Non-Infectious Molecular Clone NL4-3 Gag-iGFP ΔEnv (ARP-12455), contributed by Dr. Benjamin K. Chen; Anti-HIV-1 gp120 Monoclonal Antibody PGT128 (ARP-13352), contributed by the International AIDS Vaccine Initiative; HIV-1 SG3ΔEnv (ARP-11051), contributed by Dr. John C. Kappes and Dr. Christina Ochsenbauer; and HIV-1 Gag-iGFP JRFL (ARP-12456), contributed by Dr. Benjamin K. Chen. We thank BioLegend for providing the CD38 and CD44 PE-labeled antibodies. During the preparation of this work, JB used ChatGPT version 5.2 in order to edit and enhance readability of the text. After using this tool, the authors reviewed and edited the content and take full responsibility for the content.

## References

1. Fernandes C, Persaud AT, Chaphekhar D, et al. Flow virometry: recent advancements, best practices, and future frontiers. J Virol. 2025;99(2):e01717–24. doi:10.1128/jvi.01717-24

2. Zamora JLR, Aguilar HC. Flow virometry as a tool to study viruses. Methods. 2018;134-135:87–97. doi:10.1016/j.ymeth.2017.12.011

3. Lippé R. Flow Virometry: a Powerful Tool To Functionally Characterize Viruses. J Virol. 2018;92(3):10.1128/jvi.01765-17. doi:10.1128/jvi.01765-17

4. Tabler CO, Tilton JC. Analysis of Individual Viral Particles by Flow Virometry. Viruses. 2024;16(5):802. doi:10.3390/v16050802

5. Hussain R, Ongaro AE, Concepción MLR de la, et al. Small form factor flow virometer for SARS-CoV-2. Biomed Opt Express. 2022;13(3):1609–1619. doi:10.1364/BOE.450212

6. Soni N, Pai P, Krishna Kumar GR, Prasad V, Dasgupta S, Bhadra B. A Flow Virometry Process Proposed for Detection of SARS-CoV-2 and large-scale Screening of COVID-19 Cases. Future Virol. 2020;15(8):525–532. doi:10.2217/fvl-2020-0141

7. Samman N, El-Boubbou K, Al-Muhalhil K, et al. MICaFVi: A Novel Magnetic Immuno-Capture Flow Virometry Nano-Based Diagnostic Tool for Detection of Coronaviruses. Biosensors. 2023;13(5):553. doi:10.3390/bios13050553

8. Burnie J, Persaud AT, Thaya L, et al. P-selectin glycoprotein ligand-1 (PSGL-1/CD162) is incorporated into clinical HIV-1 isolates and can mediate virus capture and subsequent transfer to permissive cells. Retrovirology. 2022;19(1):9. doi:10.1186/s12977-022-00593-5

9. El Bilali N, Duron J, Gingras D, Lippé R. Quantitative Evaluation of Protein Heterogeneity within Herpes Simplex Virus 1 Particles. J Virol. 2017;91(10):e00320–17. doi:10.1128/JVI.00320-17

10. Yi S, McCracken R, Davide J, et al. Development of process analytical tools for rapid monitoring of live virus vaccines in manufacturing. Sci Rep. 2022;12(1):15494. doi:10.1038/s41598-022-19744-x

11. Ricci G, Minsker K, Kapish A, et al. Flow virometry for process monitoring of live virus vaccineslessons learned from ERVEBO. Sci Rep. 2021;11(1):7432. doi:10.1038/s41598-021-86688-z

12. Hercher M, Mueller W, Shapiro HM. Detection and discrimination of individual viruses by flow cytometry. J Histochem Cytochem. 1979;27(1):350–352. doi:10.1177/27.1.374599

13. Gaudin R, Barteneva NS. Sorting of small infectious virus particles by flow virometry reveals distinct infectivity profiles. Nat Commun. 2015;6:6022. doi:10.1038/ncomms7022

14. Brussaard CPD. Optimization of Procedures for Counting Viruses by Flow Cytometry. Appl Environ Microbiol. 2004;70(3):1506–1513. doi:10.1128/AEM.70.3.1506-1513.2004

15. Marie D, Brussaard CPD, Thyrhaug R, Bratbak G, Vaulot D. Enumeration of Marine Viruses in Culture and Natural Samples by Flow Cytometry. APPL Env MICROBIOL. 1999;65:8.

16. Brussaard CPD, Marie D, Bratbak G. Flow cytometric detection of viruses. J Virol Methods. 2000;85(1):175–182. doi:10.1016/S0166-0934(99)00167-6

17. Khalil JYB, Langlois T, Andreani J, et al. Flow Cytometry Sorting to Separate Viable Giant Viruses from Amoeba Co-culture Supernatants. Front Cell Infect Microbiol. 2017;6:202. doi:10.3389/fcimb.2016.00202

18. Khalil JYB, Robert S, Reteno DG, Andreani J, Raoult D, La Scola B. High-Throughput Isolation of Giant Viruses in Liquid Medium Using Automated Flow Cytometry and Fluorescence Staining. Front Microbiol. 2016;7:26. doi:10.3389/fmicb.2016.00026

19. Khadivjam B, El Bilali N, Lippé R. Analysis and Sorting of Individual HSV-1 Particles by Flow Virometry. In: Diefenbach RJ, Fraefel C, eds. Herpes Simplex Virus : Methods and Protocols. Springer; 2020:289–303. doi:10.1007/978-1-4939-9814-2_16

20. Loret S, El Bilali N, Lippé R. Analysis of herpes simplex virus type I nuclear particles by flow cytometry. Cytometry A. 2012;81A(11):950–959. doi:10.1002/cyto.a.22107

21. Bonar MM, Tilton JC. High sensitivity detection and sorting of infectious human immunodeficiency virus (HIV-1) particles by flow virometry. Virology. 2017;505:80–90. doi:10.1016/j.virol.2017.02.016

22. Bonar MM, Tabler CO, Haqqani AA, et al. Nanoscale flow cytometry reveals interpatient variability in HIV protease activity that correlates with viral infectivity and identifies drug-resistant viruses. Sci Rep. 2020;10(1):18101. doi:10.1038/s41598-020-75118-1

23. Arakelyan A, Fitzgerald W, Margolis L, Grivel JC. Nanoparticle-based flow virometry for the analysis of individual virions. J Clin Invest. 2013;123(9):3716–3727. doi:10.1172/JCI67042

24. Zicari S, Arakelyan A, Fitzgerald W, et al. Evaluation of the maturation of individual Dengue virions with flow virometry. Virology. 2016;488:20–27. doi:10.1016/j.virol.2015.10.021

25. Arakelyan A, Petersen JD, Blazkova J, Margolis L. Macrophage-derived HIV-1 carries bioactive TGF-beta. Sci Rep. 2019;9(1):1. doi:10.1038/s41598-019-55615-8

26. Yan X, Schielke EG, Grace KM, Hassell C, Marrone BL, Nolan JP. Microsphere-based duplexed immunoassay for influenza virus typing by flow cytometry. J Immunol Methods. 2004;284(1):27–38. doi:10.1016/j.jim.2003.09.016

27. Samman N, Aljami HA, Alhayli S, et al. Implementation and validation of MICaFVi: A highly efficient nanotechnology-based method for coronaviruses detection. Sens Actuators Rep. 2024;8:100248. doi:10.1016/j.snr.2024.100248

28. Arakelyan A, Fitzgerald W, Zicari S, Vanpouille C, Margolis L. Extracellular Vesicles Carry HIV Env and Facilitate Hiv Infection of Human Lymphoid Tissue. Sci Rep. 2017;7:1695. doi:10.1038/s41598-017-01739-8

29. Brittain GC, Chen YQ, Martinez E, et al. A Novel Semiconductor-Based Flow Cytometer with Enhanced Light-Scatter Sensitivity for the Analysis of Biological Nanoparticles. Sci Rep. 2019;9:16039. doi:10.1038/s41598-019-52366-4

30. Morales-Kastresana A, Telford B, Musich TA, et al. Labeling extracellular vesicles for nanoscale flow cytometry. Sci Rep. 2017;7(1):1878.

31. Morales-Kastresana A, Musich TA, Welsh JA, et al. High-fidelity detection and sorting of nanoscale vesicles in viral disease and cancer. J Extracell Vesicles. 2019;8(1):1597603. doi:10.1080/20013078.2019.1597603

32. Sharma M, Sheth M, Poling HM, Kuhnell D, Langevin SM, Esfandiari L. Rapid purification and multiparametric characterization of circulating small extracellular vesicles utilizing a label-free lab-on-a-chip device. Sci Rep. 2023;13(1):1. doi:10.1038/s41598-023-45409-4

33. O’Leary M L., Bowtell J, Richards M, et al. Effects of the DailyColors™ polyphenol supplement on serum proteome, cognitive function, and health in older adults at risk of cognitive and functional decline. Food Funct. Published online 2025. doi:10.1039/D4FO06259K

34. Cook S, Tang VA, Lannigan J, Jones JC, Welsh JA. Quantitative flow cytometry enables end-to-end optimization of cross-platform extracellular vesicle studies. Cell Rep Methods. 2023;3(12). doi:10.1016/j.crmeth.2023.100664

35. Renner TM, Tang VA, Burger D, Langlois MA. Intact Viral Particle Counts Measured by Flow Virometry Provide Insight into the Infectivity and Genome Packaging Efficiency of Moloney Murine Leukemia Virus. J Virol. 2020;94(2):10.1128/jvi.01600-19. doi:10.1128/jvi.01600-19

36. Welsh JA, Jones JC, Tang VA. Fluorescence and Light Scatter Calibration Allow Comparisons of Small Particle Data in Standard Units across Different Flow Cytometry Platforms and Detector Settings. Cytometry A. 2020;97(6):592–601. doi:10.1002/cyto.a.24029

37. Maltseva M, Langlois MA. Influence of GlycoGag on the Incorporation of Host Membrane Proteins Into the Envelope of the Moloney Murine Leukemia Virus. Front Virol. 2021;1. doi:10.3389/fviro.2021.747253

38. Tang VA, Fritzsche AK, Renner TM, et al. Engineered Retroviruses as Fluorescent Biological Reference Particles for Small Particle Flow Cytometry. bioRxiv. Preprint posted online June 7, 2019:614461. doi:10.1101/614461

39. Maltseva M, Langlois MA. Flow Virometry for Characterizing the Size, Concentration, and Surface Antigens of Viruses. Curr Protoc. 2022;2(2):e368. doi:10.1002/cpz1.368

40. Burnie J, Tang VA, Welsh JA, et al. Flow Virometry Quantification of Host Proteins on the Surface of HIV-1 Pseudovirus Particles. Viruses. 2020;12(11):11. doi:10.3390/v12111296

41. Persaud AT, Khela J, Fernandes C, et al. Virion-incorporated CD14 enables HIV-1 to bind LPS and initiate TLR4 signaling in immune cells. J Virol. 98(5):e00363–24. doi:10.1128/jvi.00363-24

42. Burnie J, Fernandes C, Patel A, et al. Applying Flow Virometry to Study the HIV Envelope Glycoprotein and Differences Across HIV Model Systems. Viruses. 2024;16(6):6. doi:10.3390/v16060935

43. Chaphekar D, Fernandes C, Persaud AT, Guzzo C. Comparing methods to detect cellular proteins on the surface of HIV-1 virions. J Virol Methods. Published online December 6, 2024:115096. doi:10.1016/j.jviromet.2024.115096

44. Burnie J, Fernandes C, Chaphekar D, et al. Identification of CD38, CD97, and CD278 on the HIV surface using a novel flow virometry screening assay. Sci Rep. 2023;13(1):23025. doi:10.1038/s41598-023-50365-0

45. Maltseva M, Rossotti MA, Tanha J, Langlois MA. Characterization of Nanobody Binding to Distinct Regions of the SARS-CoV-2 Spike Protein by Flow Virometry. Viruses. 2025;17(4):4. doi:10.3390/v17040571

46. Application Note: CytoFLEX nano Flow Cytometer: the new frontier of nanoscale Flow Cytometry. Accessed January 22, 2025. https://www.nature.com/articles/d42473-024-00117-z

47. Lin YH, Chen Y, Liu EW, et al. Immunomodulation effects of collagen hydrogel encapsulating extracellular vesicles derived from calcium silicate stimulated-adipose mesenchymal stem cells for diabetic healing. J Nanobiotechnology. 2025;23(1):45. doi:10.1186/s12951-025-03097-4

48. Chen YW, Lin YH, Ho CC, et al. High-yield extracellular vesicle production from HEK293T cells encapsulated in 3D auxetic scaffolds with cyclic mechanical stimulation for effective drug carrier systems. Biofabrication. 2024;16(4):045035. doi:10.1088/1758-5090/ad728b

49. Gudbergsson JM, Malle MG, Jensen JB, et al. Dedicated nanoparticle flow cytometry for single extracellular vesicle phenotyping: Performance of the CytoFLEX Nano. bioRxiv. Preprint posted online December 17, 2025:2025.12.15.694348. doi:10.64898/2025.12.15.694348

50. Cao L, Wang H, Zhu J, et al. Exploring exosome profiling via CytoFLEX Nano flow cytometer: Approaches and applications. VIEW. 2025;6(6):20250068. doi:10.1002/VIW.20250068

51. Bokun V, Strang BL, Pantazi P, Liu Y, Holder B. Nano-Flow Cytometry-Guided Discrimination and Separation of Human Cytomegalovirus Virions and Extracellular Vesicles. J Extracell Vesicles. 2025;14(5):e70060. doi:10.1002/jev2.70060

52. Nolte-’t Hoen E, Cremer T, Gallo RC, Margolis LB. Extracellular vesicles and viruses: Are they close relativeš Proc Natl Acad Sci. 2016;113(33):9155–9161. doi:10.1073/pnas.1605146113

53. van der Grein SG, Defourny KAY, Slot EFJ, Nolte-’t Hoen ENM. Intricate relationships between naked viruses and extracellular vesicles in the crosstalk between pathogen and host. Semin Immunopathol. 2018;40(5):491–504. doi:10.1007/s00281-018-0678-9

54. Welsh JA, Jones JC. Small Particle Fluorescence and Light Scatter Calibration Using FCMPASS Software. Curr Protoc Cytom. 2020;94(1):e79. doi:10.1002/cpcy.79

55. Welsh JA, Arkesteijn GJA, Bremer M, et al. A compendium of single extracellular vesicle flow cytometry. J Extracell Vesicles. 2023;12(2):e12299. doi:10.1002/jev2.12299

56. Welsh JA, Van Der Pol E, Arkesteijn GJA, et al. MIFlowCyt-EV: a framework for standardized reporting of extracellular vesicle flow cytometry experiments. J Extracell Vesicles. 9(1):1713526. doi:10.1080/20013078.2020.1713526

57. Arce JE, Welsh JA, Cook S, et al. The NanoFlow Repository. Bioinformatics. 2023;39(6):btad368. doi:10.1093/bioinformatics/btad368

58. Burnie J, Guzzo C. The Incorporation of Host Proteins into the External HIV-1 Envelope. Viruses. 2019;11(1):85. doi:10.3390/v11010085

59. Tremblay MJ, Fortin JF, Cantin R. The acquisition of host-encoded proteins by nascent HIV-1. Immunol Today. 1998;19(8):346–351. doi:10.1016/S0167-5699(98)01286-9

60. Orentas RJ, Hildreth JEK. Association of Host Cell Surface Adhesion Receptors and Other Membrane Proteins with HIV and SIV. AIDS Res Hum Retroviruses. 1993;9(11):1157–1165. doi:10.1089/aid.1993.9.1157

61. Cantin R, Diou J, Bélanger D, Tremblay AM, Gilbert C. Discrimination between exosomes and HIV-1: Purification of both vesicles from cell-free supernatants. J Immunol Methods. 2008;338(1):21–30. doi:10.1016/j.jim.2008.07.007

62. Moulin C, Crupi MJF, Ilkow CS, Bell JC, Boulton S. Extracellular Vesicles and Viruses: Two Intertwined Entities. Int J Mol Sci. 2023;24(2):2. doi:10.3390/ijms24021036

63. Gould SJ, Booth AM, Hildreth JEK. The Trojan exosome hypothesis. Proc Natl Acad Sci. 2003;100(19):10592–10597. doi:10.1073/pnas.1831413100

64. Hemler ME. Tetraspanin functions and associated microdomains. Nat Rev Mol Cell Biol. 2005;6(10):801–811. doi:10.1038/nrm1736

65. Charrin S, Jouannet S, Boucheix C, Rubinstein E. Tetraspanins at a glance. J Cell Sci. 2014;127(17):3641–3648. doi:10.1242/jcs.154906

66. Rubinstein E, Théry C, Zimmermann P. Tetraspanins affect membrane structures and the trafficking of molecular partners: what impact on extracellular vesicleš Biochem Soc Trans. 2025;53(02):371–382. doi:10.1042/BST20240523

67. Willms E, Cabañas C, Mäger I, Wood MJA, Vader P. Extracellular Vesicle Heterogeneity: Subpopulations, Isolation Techniques, and Diverse Functions in Cancer Progression. Front Immunol. 2018;9. doi:10.3389/fimmu.2018.00738

68. Jankovičová J, Sečová P, Michalková K, Antalíková J. Tetraspanins, More than Markers of Extracellular Vesicles in Reproduction. Int J Mol Sci. 2020;21(20):7568. doi:10.3390/ijms21207568

69. Lehrich BM, Liang Y, Fiandaca MS. Foetal bovine serum influence on in vitro extracellular vesicle analyses. J Extracell Vesicles. 2021;10(3):e12061. doi:10.1002/jev2.12061

70. Urzì O, Olofsson Bagge R, Crescitelli R. The dark side of foetal bovine serum in extracellular vesicle studies. J Extracell Vesicles. 2022;11(10):e12271. doi:10.1002/jev2.12271

71. Jolly C, Sattentau QJ. Human Immunodeficiency Virus Type 1 Assembly, Budding, and Cell-Cell Spread in T Cells Take Place in Tetraspanin-Enriched Plasma Membrane Domains. J Virol. 2007;81(15):7873–7884. doi:10.1128/JVI.01845-06

72. Ruiz-Mateos E, Pelchen-Matthews A, Deneka M, Marsh M. CD63 Is Not Required for Production of Infectious Human Immunodeficiency Virus Type 1 in Human Macrophages. J Virol. 2008;82(10):4751–4761. doi:10.1128/jvi.02320-07

73. Hübner W, Chen P, Portillo AD, Liu Y, Gordon RE, Chen BK. Sequence of Human Immunodeficiency Virus Type 1 (HIV-1) Gag Localization and Oligomerization Monitored with Live Confocal Imaging of a Replication-Competent, Fluorescently Tagged HIV-1. J Virol. 2007;81(22):12596–12607. doi:10.1128/JVI.01088-07

74. Pejchal R, Doores KJ, Walker LM, et al. A potent and broad neutralizing antibody recognizes and penetrates the HIV glycan shield. Science. 2011;334(6059):1097–1103. doi:10.1126/science.1213256

75. Bello-Morales R, Ripa I, López-Guerrero JA. Extracellular Vesicles in Viral Spread and Antiviral Response. Viruses. 2020;12(6):623. doi:10.3390/v12060623

76. DeMarino C, Denniss J, Cowen M, et al. HIV-1 RNA in extracellular vesicles is associated with neurocognitive outcomes. Nat Commun. 2024;15(1):4391. doi:10.1038/s41467-024-48644-z

77. Giannessi F, Aiello A, Franchi F, Percario ZA, Affabris E. The Role of Extracellular Vesicles as Allies of HIV, HCV and SARS Viruses. Viruses. 2020;12(5):571. doi:10.3390/v12050571

78. Schwartz A, Wang L, Early E, et al. Quantitating Fluorescence Intensity from Fluorophore: The Definition of MESF Assignment. J Res Natl Inst Stand Technol. 2002;107(1):83–91. doi:10.6028/jres.107.009

79. Wang L, Gaigalas AK, Abbasi F, Marti GE, Vogt RF, Schwartz A. Quantitating Fluorescence Intensity From Fluorophores: Practical Use of MESF Values. J Res Natl Inst Stand Technol. 2002;107(4):339–353. doi:10.6028/jres.107.027

80. Guzzo C, Ichikawa D, Park C, et al. Virion incorporation of integrin α4β7 facilitates HIV-1 infection and intestinal homing. Sci Immunol. 2017;2(11):eaam7341. doi:10.1126/sciimmunol.aam7341

81. van Niel G, D’Angelo G, Raposo G. Shedding light on the cell biology of extracellular vesicles. Nat Rev Mol Cell Biol. 2018;19(4):213–228. doi:10.1038/nrm.2017.125

82. Doyle LM, Wang MZ. Overview of Extracellular Vesicles, Their Origin, Composition, Purpose, and Methods for Exosome Isolation and Analysis. Cells. 2019;8(7):727. doi:10.3390/cells8070727

83. Sheta M, Taha EA, Lu Y, Eguchi T. Extracellular Vesicles: New Classification and Tumor Immunosuppression. Biology. 2023;12(1):110. doi:10.3390/biology12010110

84. Finzi D, Plaeger SF, Dieffenbach CW. Defective Virus Drives Human Immunodeficiency Virus Infection, Persistence, and Pathogenesis. Clin Vaccine Immunol. 2006;13(7):715–721. doi:10.1128/CVI.00052-06

