## Supplementary Figures for "Evaluating the Utility of a Nanoscale Flow Cytometer for Detection of Surface Proteins on HIV and Extracellular Vesicles"

**Fig S1. PE antibody titrations for host protein labeling.**

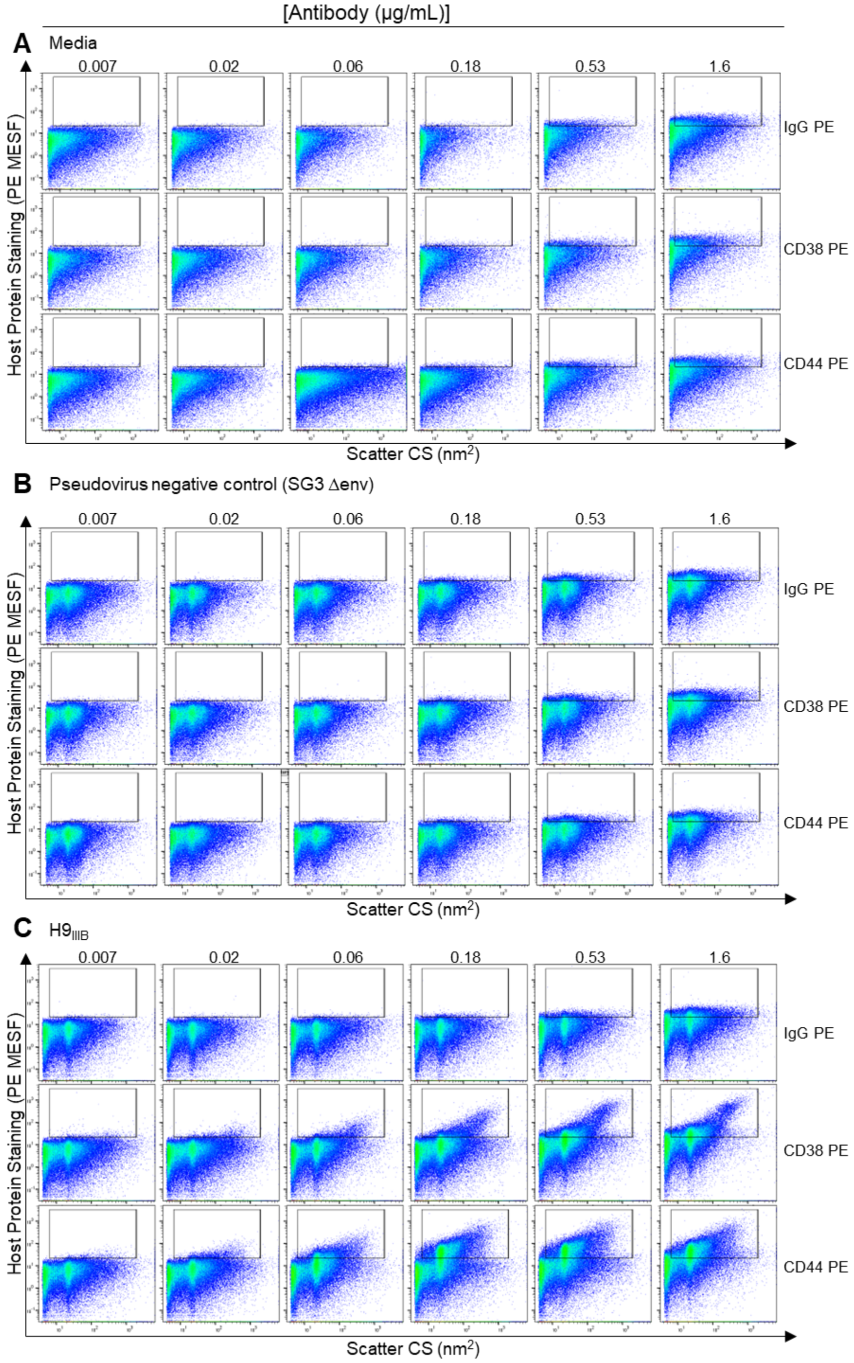


**Fig S1. PE antibody titrations for host protein labeling.** (A) PE-conjugated antibodies targeting CD38 and CD44 were titrated on cell culture media samples, (B) pseudovirus controls negative for the proteins of interest (SG3 ΔEnv), and (C) the $HIV_{IIIB}$ isolate grown in H9 cells ($H9_{IIIB}$) using the CytoFLEX S. An isotype control (IgG) is shown to assess signal from nonspecific antibody binding, with gates displaying positive staining as set on the isotype and media controls.

### Fig S2. BV421 antibody titrations for host protein labeling.

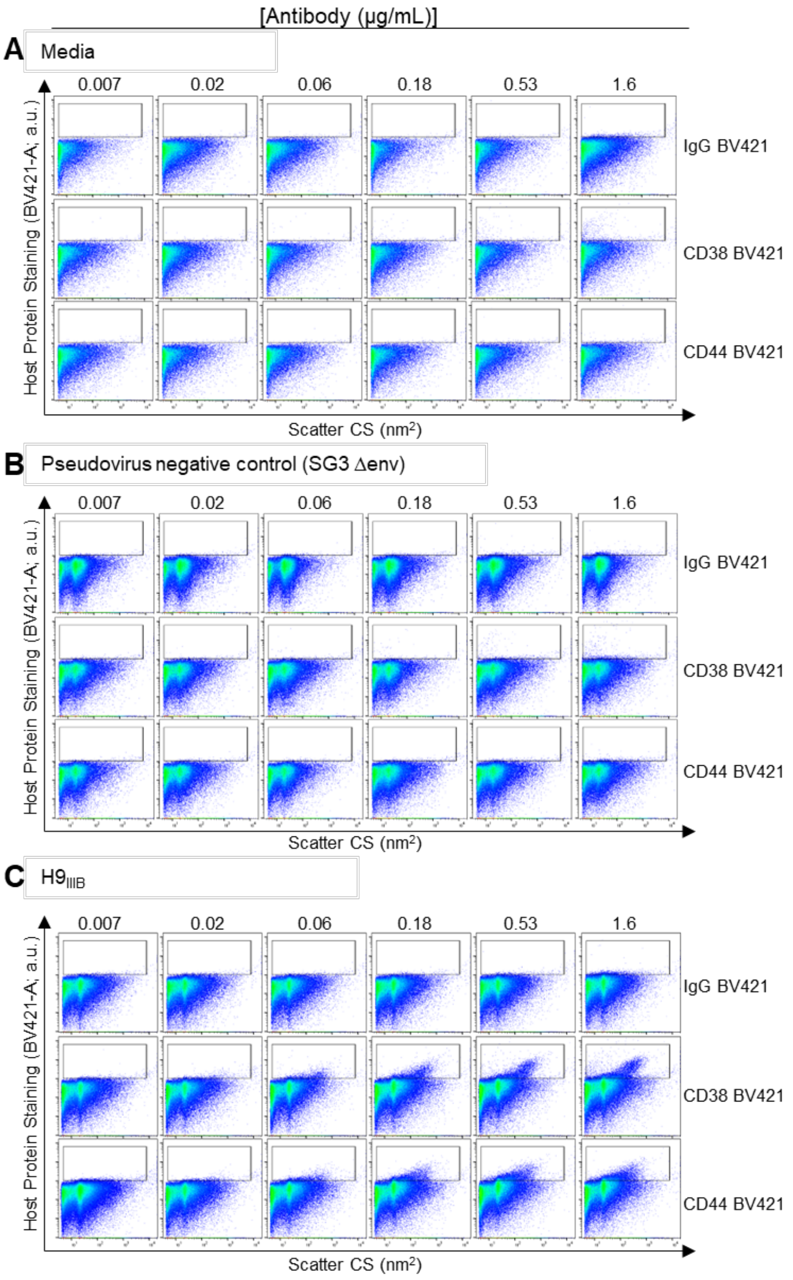


**Fig S2. BV421 antibody titrations for host protein labeling.** BV421-conjugated antibodies targeting CD38 and CD44 were titrated on cell culture media samples, (B) pseudovirus controls negative for the proteins of interest (SG3 ΔEnv), and (C) the $HIV_{IIIB}$ isolate using CytoFLEX S. An isotype control (IgG) is shown to assess signal from nonspecific antibody binding, with gates displaying positive staining as set on the isotype and media controls.

### Fig S3. BV421 antibody titrations for tetraspanin staining.

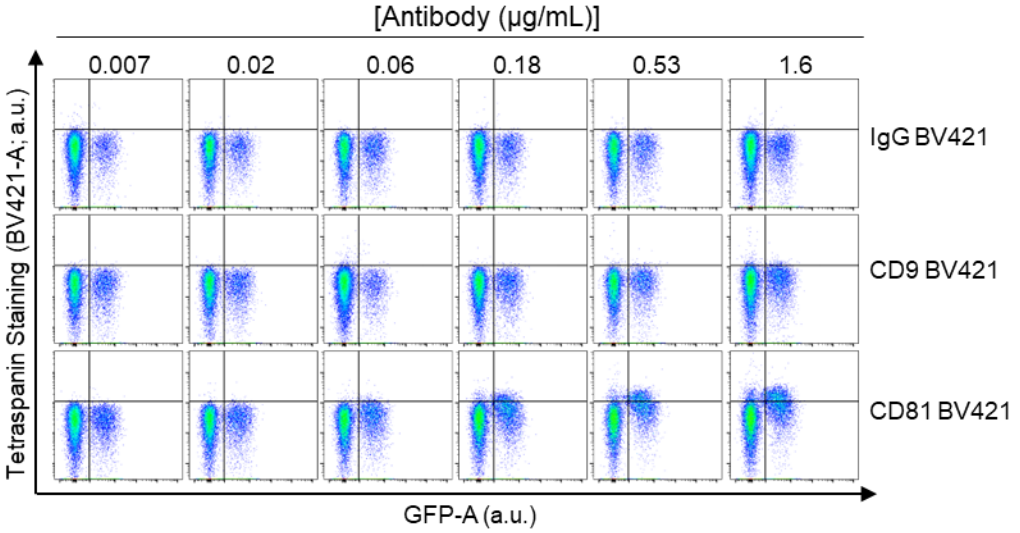


**Fig S3. BV421 antibody titrations for tetraspanin staining.**
Titrations of BV421-conjugated antibodies against CD9, CD63, and CD81 cellular proteins on HIV iGFP JR FL virus tested on CytoFLEX S. An isotype control (IgG) is shown to assess background levels of fluorescence. Gates are set on the isotype control.

### Fig S4. Antibody titration and controls for indirect Env labeling.

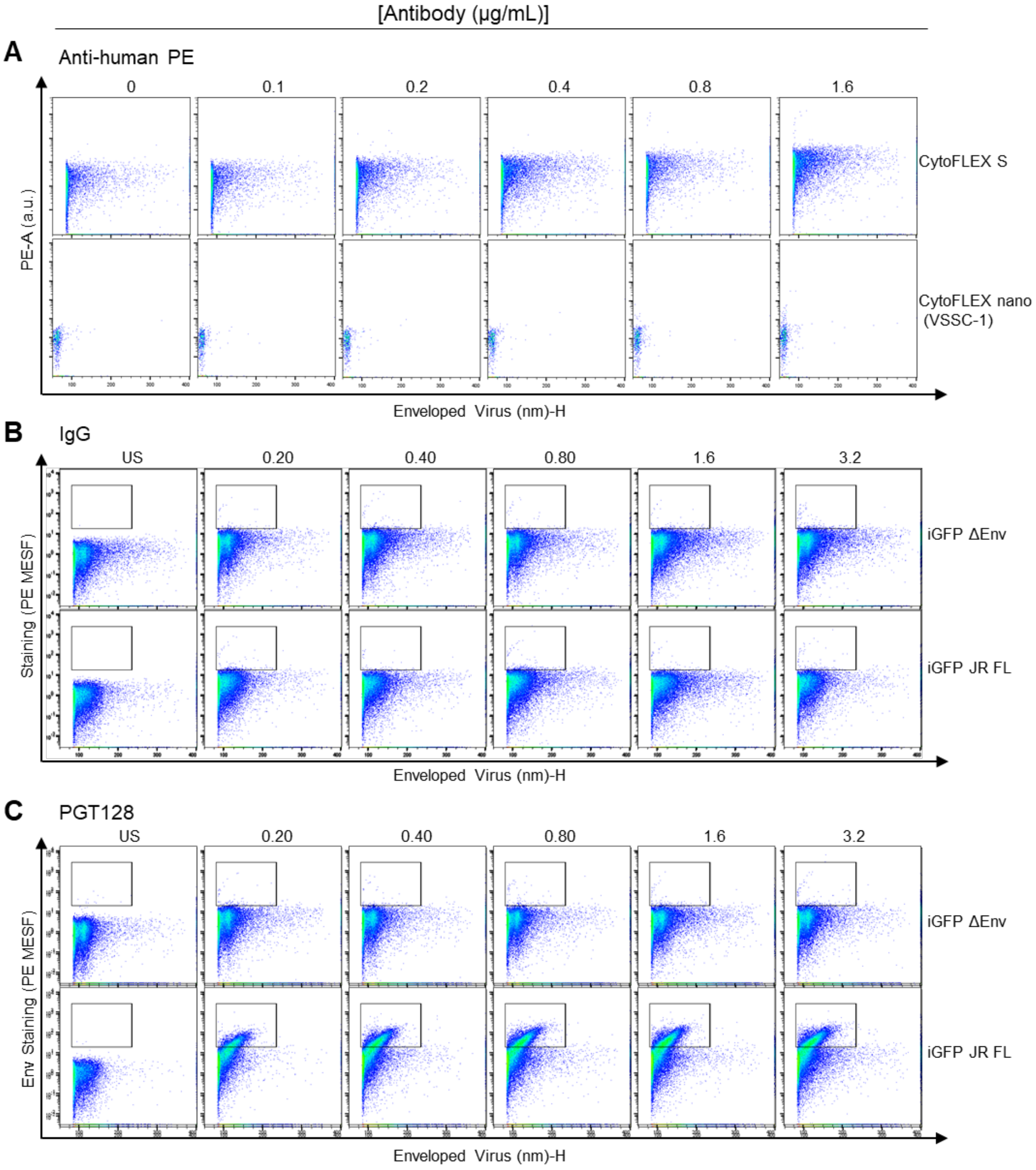


**Fig S4. Antibody titration and controls for indirect Env labeling.** (A) Titration of PE-conjugated anti-human secondary antibody to distinguish background fluorescent noise from secondary antibody alone on CytoFLEX S (top panels) and CytoFLEX nano (bottom panels). (B) Titration of isotype control (IgG) or anti-Env antibody, (C) PGT128 tested on HIV iGFP virions with (JR FL) or without (ΔEnv) the HIV envelope glycoprotein on the CytoFLEX S to optimize signal-to-noise ratios.

### Fig S5. Host protein staining on the CytoFLEX nano with acquisition by time.

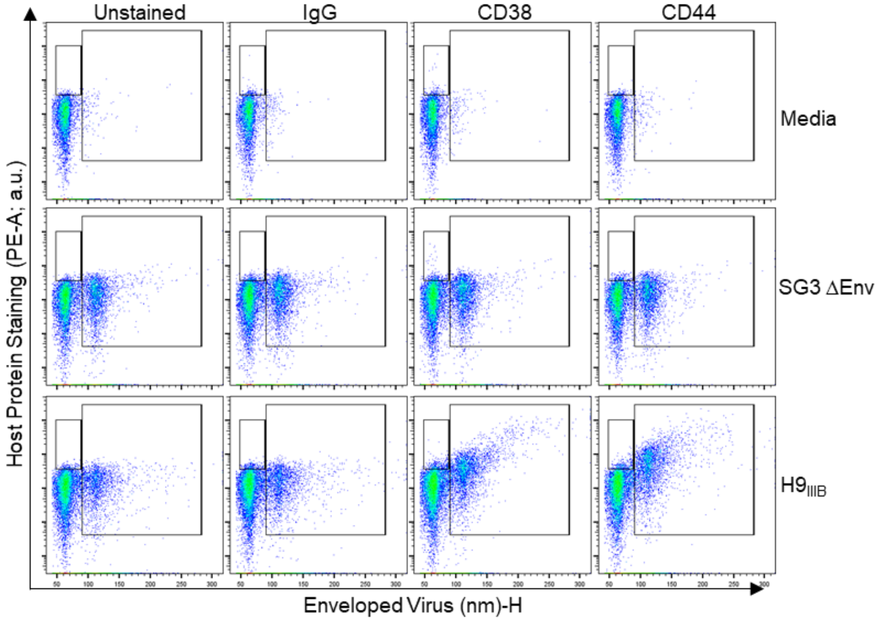


**Figure S5. Host protein staining on the CytoFLEX nano with acquisition by time.** Samples were stained as in Fig. 2 and acquired for 15 seconds. The right gate is set based on the virus population as determined by scatter and is inclusive of the entire stained population. The left gate is set above instrument background fluorescence at the instrument threshold and includes smaller particles including EVs and unbound antibody.

**Fig S6. Serial dilutions of virus samples.**

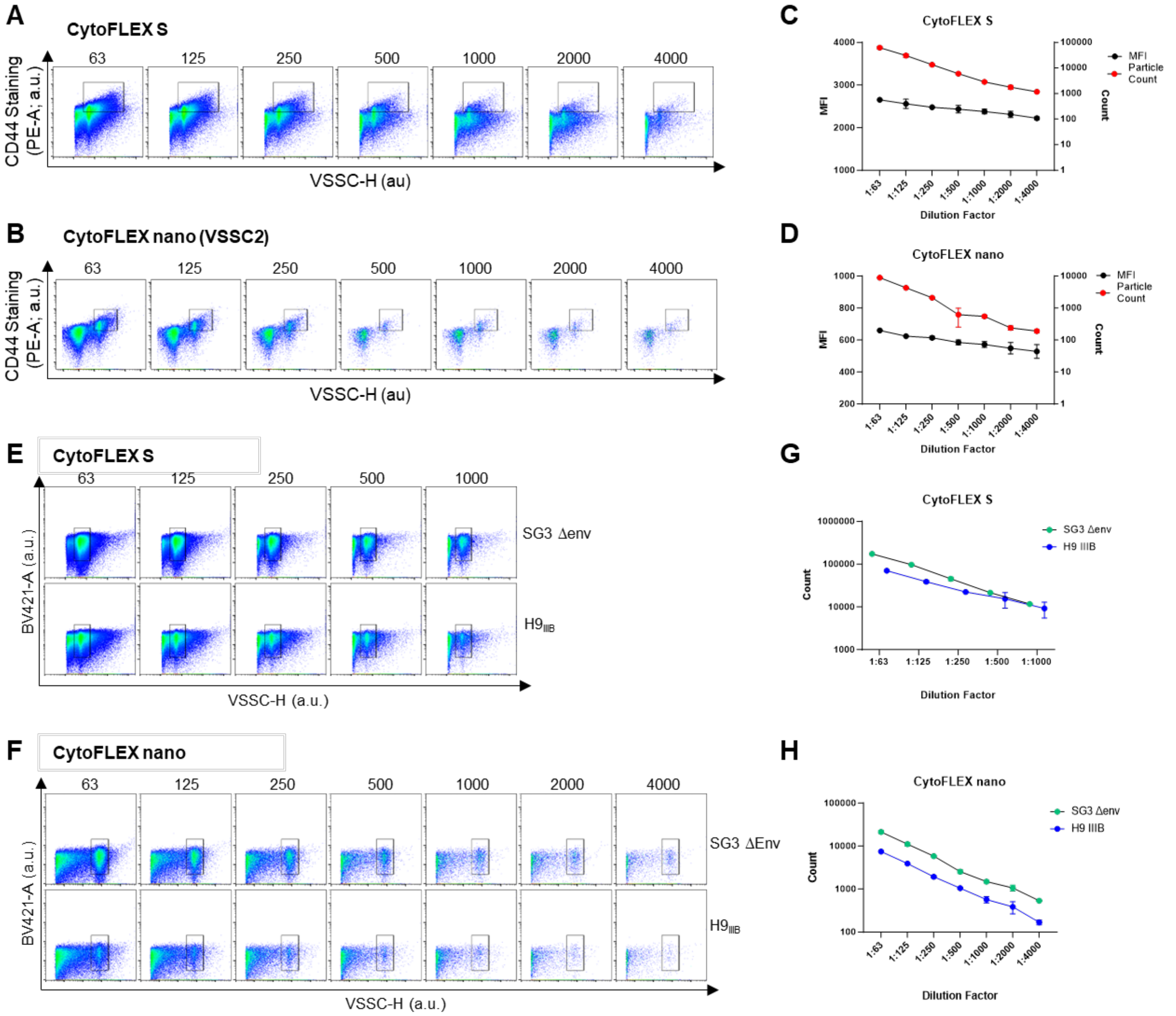


**Fig S6. Serial dilutions of virus samples.** (A) Serial dilutions of CD44 PE-stained $H9_{IIIB}$ were analyzed on the CytoFLEX S and (B) CytoFLEX nano as flow virometry controls. (C, D) Quantitation of event counts and median fluorescence intensity from the gated regions shown in A and B. (E) Serial dilutions of unstained replication-competent HIV ($H9_{IIIB}$; top panels) and HIV pseudoviruses (SG3 ΔEnv; bottom panels) were analyzed on the CytoFLEX S and (F) CytoFLEX nano. (G, H) Quantitation of event counts from the gated regions shown in E and F, respectively.

### Fig S7. Dual labeling media controls.

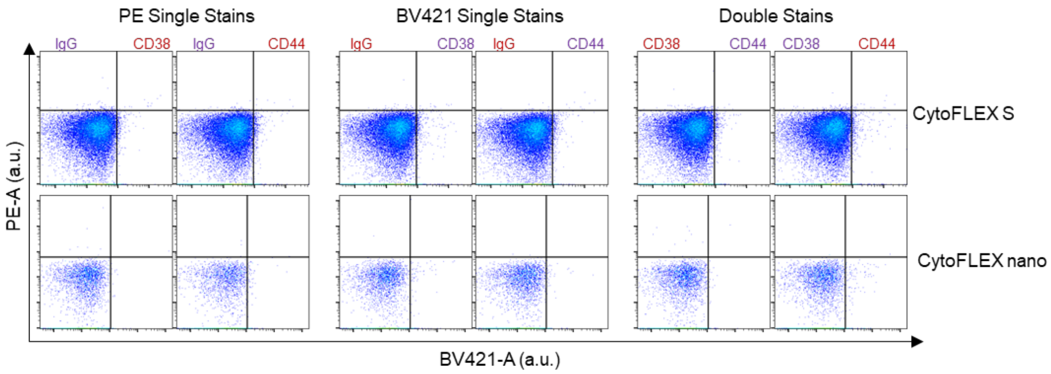


**Fig S7. Dual labeling media controls.** (A) Cell culture media was stained using anti-CD38 and anti-CD44 antibodies on CytoFLEX S and CytoFLEX nano cytometers. Red and purple labels represent PE and BV421 conjugated antibodies, respectively. Isotype controls (IgG) were used to evaluate nonspecific staining for each channel.
